# Item recognition is associated with gut microbiota composition in healthy humans

**DOI:** 10.1101/2025.04.10.648258

**Authors:** JP Oyarzun, TM Kuntz, XC Morgan, L Davachi, C Huttenhower, JE LeDoux, EA Phelps

**Author notes:** Corresponding author., 52 Oxford Street, Cambridge, MA, United States 02138.

## Abstract

Murine studies show that the gut microbiota – the collection of the microbes residing in the large intestine – affects memory performance in the host. However, whether commensal gut bacteria are linked to human episodic memory remains unknown. Here, we investigated whether individual differences in episodic memory performance were associated with differences in the indigenous gut microbiota composition between individuals. We show that greater gut microbiota alpha diversity was associated with better item recognition and that gut microbiota dissimilarity index (beta diversity) between participants was associated with differences in their performance. Finally, our results suggest that *Prevotella copri* might play a role in the relationship between gut microbiota and human item recognition in healthy individuals. In a sample size larger than previous human studies and examining unmanipulated gut microbiota, we provide evidence that episodic memory in healthy humans is linked to their gut microbiota composition.

**SIGNIFICANCE STATEMENT:** Understanding the relationship between the gut microbiota – the collective of microbes in our gut – and human cognition represents a pivotal frontier in neuroscience. Our study dives into this quest by exploring the connection between the indigenous gut microbiota and episodic memory in healthy individuals. While prior research has illuminated this relationship in murine models, our study extends these findings to humans, providing compelling evidence that gut microbiota diversity and structural composition are associated with differences in episodic memory performance. By shifting the focus from probiotic-based interventions to the study of our commensal gut microbes, our findings shed light on how the gut microbiota, in its natural state, could be linked to interindividual differences in human cognition.

## INTRODUCTION

The ability to form memories is a fundamental cognitive mechanism that allows individuals to predict their environment and adapt their behavior accordingly. Memory malfunction contributes to several disorders, including depression, anxiety, and dementia^1^. For this reason, identifying modifiable factors that affect memory function could reveal new treatment targets.

Recent evidence in rodent models has shown that the gut microbiota, the collection of the microbes residing in the large intestine, affects memory performance and hippocampal structure and function in the host ^2–4^. For example, while the absence of gut microbes in germ-free and antibiotic-treated rodents impairs memory recognition^2,5^, administering certain live bacteria species, called probiotics, can improve it^6^. In turn, administration of an enteric pathogenic bacteria can impair memory recognition in germ-free or stressed mice, but performance can be preserved if animals receive preventive treatment with probiotics^7^. Similarly, studies employing a “cafeteria diet” or “Western diet” show a significant impairment of memory recognition and spatial learning in rats^8– 10^. This work demonstrates that such high-sugar, high-fat diets reduce the diversity and alter the gut microbiota composition, which is linked to memory impairment^8,10,11^. Again, preventive treatment with probiotics can prevent diet-induced memory impairments^12^.

Unlike findings in rodents, evidence linking gut microbiota composition and memory performance in humans is less clear. Most studies have treated healthy individuals for 3 to 12 weeks with one or more probiotics, after which they compared differences in performance either pre- and post-treatment or between control and placebo groups. For example, a study of 22 males using a within-subject design showed that compared to pre-treatment baseline, 4-week administration of *Bifidobacterium longum* resulted in a greater increase in performance - in a paired associated memory task - than after placebo treatment. However, a direct comparison of memory performance between probiotic and placebo groups following treatment was not significant^13^. In a similarly designed study with 45 participants using a multi-species probiotic, the treated group showed better memory recognition for unpleasant emotional pictures but not neutral ones^14^. In contrast, a larger sample of 120 older adults (average age of 61.8), using a 10-day treatment with fermented milk containing *Lactobacillus casei* (strain Shirota), showed a decrease in episodic memory performance compared to a placebo group (using the Wechsler Memory Scale)^15^. Another study with a 12-week treatment and randomized, double-blind, and placebo-controlled design showed improved verbal learning and memory in stressed participants treated with *Lactobacillus plantarum*. Still, this effect was only marginally significant in men and non-significant in women^16^. In a similar design, 32 older adults (age 60-75) treated with *Lactobacillus helveticus* administered via fermented milk significantly improved memory and attention compared to a placebo group^17^. Lastly, Alzheimer patients treated with multiple-species probiotic improved their overall cognitive performance measured with the MMSE^18^.

Although these studies using probiotic manipulations suggest a link between human microbiome composition and episodic memory, the effects are often small, and the findings of a benefit are inconsistent^14^. The discrepancy between mice and humans may be due to neurobiological differences across species, lower heterogeneity in rodent model systems, and the lack of concordance between the indigenous gut microbiota profile between rodents and humans^19^. Moreover, commercial probiotics do not tend to change the microbiome composition, and their effects can differ among participants depending on their basal/ endogenous gut microbiota composition^20^. Nevertheless, given that transitory live bacteria can induce behavioral effects, we hypothesized that commensal gut species could also influence the memory system in healthy individuals. It remains unknown if the gut microbiota in its homeostatic state is related to human memory and, if this is the case, which features or taxa are relevant for memory performance.

A more direct approach to identifying relevant microbes related to human memory is to screen gut microbiota profiles in humans associated with their memory performance. Along these lines, a study with 69 older individuals (60-75 years) showed that participants’ relative abundance of microbes from the Carnobacteriaceae family was correlated with episodic memory performance^21^. However, given the small sample size and older population age, it is unclear if these results would generalize to other healthy populations.

In the current study, we investigated whether the gut microbiota composition in 127 healthy adults is associated with individual differences in episodic memory. In addition to assessing item recognition, we also examined whether more specific memory processes – pattern completion and pattern separation - were associated with microbiota composition using 16S rRNA sequencing. These processes are considered essential for the encoding and retrieval of distinct memory representations^22^. Pattern separation allows the encoding and storing of similar episodes into differentiated memory traces so we can distinctly retrieve and discriminate similar memories. Pattern completion allows individuals to retrieve memories when encountering partial or degraded retrieval cues^23^. Both memory functions are necessary for accurate, detailed recollection and have been shown to be impaired in psychiatric pathologies such as anxiety and depression^24,25^. We included three behavioral measures of memory because pattern separation and completion have been associated with distinct hippocampal structures ^23,26^. In contrast, item recognition may additionally rely on processes supported by nearby medial temporal cortical regions ^27,28^, thus allowing us to test whether these neurally distinct processes are independently associated with gut microbiota composition.

To that end, we assessed item recognition, pattern separation, and pattern completion using a mnemonic similarity task. We show that gut microbiota composition – measured as beta diversity – is associated with participants’ performance in item recognition but not independently with pattern completion or pattern separation. In addition, we found that greater alpha diversity is positively associated with better item recognition. Finally, we pinpoint a specific taxon that may contribute to the link between gut microbiota and human item recognition.

## RESULTS

### Participants showed variability in memory performance

Out of 127 enrolled participants, one hundred and fifteen answered at least 80% of trials during the incidental encoding session and were included in the analyses. Seven participants were further excluded due to incomplete metadata (n=2), low-quality sequencing data (n=4), or below-chance item recognition performance (n=1). Our final analysis included 108 participants (63 women and 45 men) with a mean age of 25.59 years (std=6.89) and a mean BMI of 24.25 (SD=4.3). Consistent with previous studies using the Mnemonic Similarity Task^29^, participants performed better at item memory recognition assessment (M= 67.72%, SD=18.95, min=7.41, max=94.76) than pattern completion (M= 31.09%, SD=15.43, min=-4.84, max=74.31) and pattern separation (M= 29.34%, SD= 19.39, min=-19.59, max=78.12) (see **Figure 1** and Methods for task details and scores calculation).

**Figure 1.**
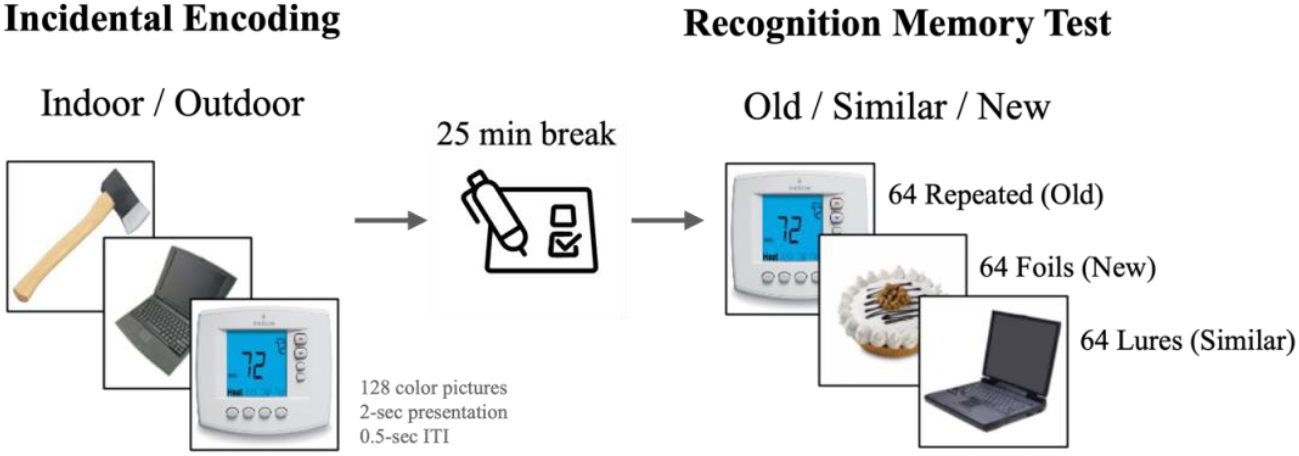
Mnemonic Similarity Task. During the incidental encoding session, participants were instructed to categorize each image as an ‘indoor’ or ‘outdoor’ object. After a 25-minute break, participants completed a surprise recognition memory test, where they viewed 64 *repetitive* items (identical to those seen in the encoding session), 64 *foil* items (entirely new objects), and 64 *lure* items (visually similar but not identical to previously seen objects). Participants were instructed to indicate if the object was ‘old’ (*repetitive*), ‘new’ (*foil*), or ‘similar’ (*lure*). *Item recognition* was calculated as the number of ‘old’ responses to *repetitive* items minus the number of ‘old’ responses to *foil* items. *Pattern Completion* was calculated as the number of ‘old’ responses to *lure* items minus the number of ‘old’ responses to *foil* items. *Pattern separation* was calculated as the number of ‘similar’ responses to *lure* items minus the number of ‘similar’ responses to *foil* items.

We taxonomically profiled participants’ fecal microbiomes (see stacked bars for species in **Table S1**) and calculated associated ecological metrics, including alpha diversity (measured as the Inverse Simpson Index) and beta diversity (measured as the Bray-Curtis distance) and single feature associations with MaAsLin2. Every model included all covariates.

### Item recognition positively correlates with alpha diversity

Alpha diversity – reflecting both richness and evenness – was positively associated with item recognition (multivariable ANOVA, F_1_= 4.04, p=0.04, **Figure 2, Table S1A**), indicating that participants with greater gut microbiota alpha diversity – a greater variety of species and a more balanced ecosystem – showed better item recognition performance. The model also showed a significant association between alpha diversity and breastfeeding, where non-breastfed individuals had higher alpha diversity, than those breastfed. No significant association was found between alpha diversity and pattern completion (F_1_= 0.05, p=0.82, **Table S1B**) or pattern separation (F_1_= 1.58, p=0.21, **Table S1C**).

**Figure 2.**
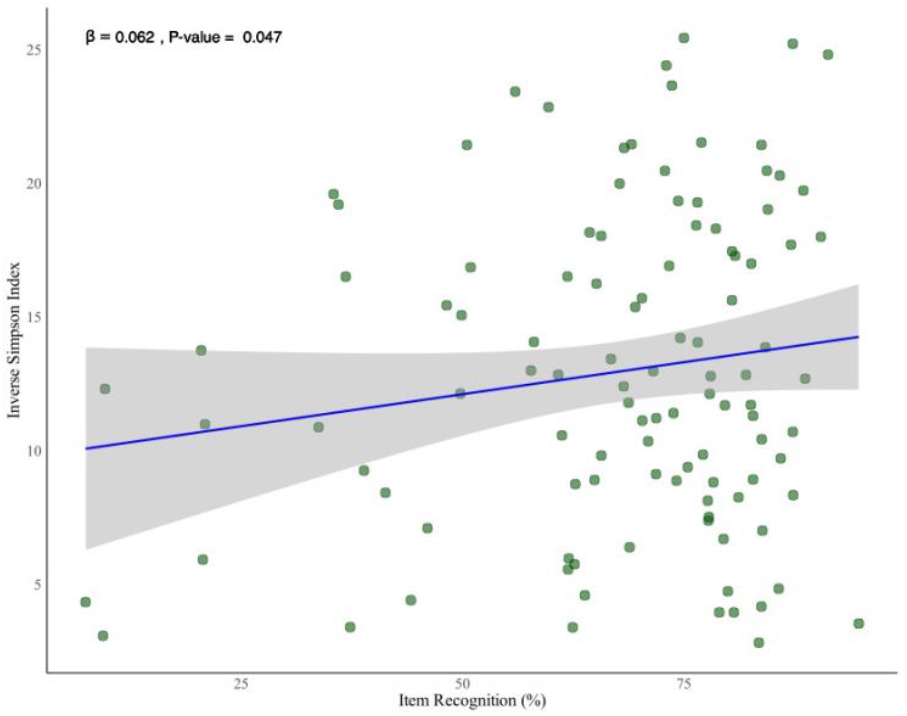
Greater alpha diversity is associated with better item recognition. A multivariable ANOVA revealed a significant relationship between alpha diversity and item recognition (*F*_1_ = 4.04, *p* = 0.04), where higher gut microbiota alpha diversity, measured by the Inverse Simpson Index, was associated with better item recognition performance.

### Item Recognition is associated with Beta Diversity

Beta diversity refers to differences in gut microbiota composition between individuals, providing insights into how unique or similar their microbial communities are. Here, we used Bray-Curtis dissimilarity to quantify these differences based on species-level abundances. Using multivariable PERMANOVA, we tested whether variation in gut microbiota composition explained differences in item recognition performance while controlling for all covariates. The significant association we observed suggests that individuals with similar gut microbiota compositions tend to have more similar item recognition performance (F_1_= 1.886, R^2^=0.017, p=0.037, **Figure 3, Table S2A**). This finding was specific to item recognition and was not observed for pattern completion (F_1_= 0.943, R^2^=0.009, p=0.466, **Table S2B**) or pattern separation (F_1_= 0.982, R^2^=0.009, p=0.411, **Table S2C**).

**Figure 3.**
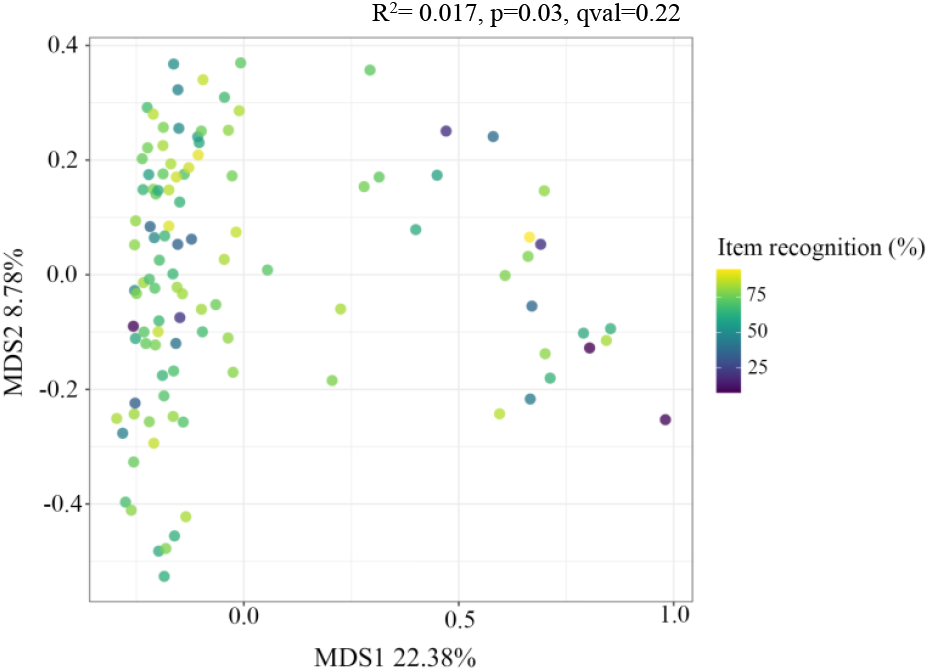
Principal coordinate analysis (PCoA) of microbiota composition using Bray-Curtis distance. PCoA plot illustrating the major axes of variation in gut microbiota composition based on Bray-Curtis distances. While item recognition was not associated with the two major axes of variation, it showed a significant association with beta diversity using a multivariable PERMANOVA on Bray-Curtis distances (R^2^ = 0.017, p = 0.03, qval = 0.22)

### Microbial features associated with item recognition

We used Microbiome Multivariable Associations with Linear Models (MaAsLin2) to test associations between specific taxa and participants’ memory performance. MaAsLin2 is specifically designed to handle sparse, non-normally distributed microbiome data, employing arcsine square root transformation to spread abundance values while preserving zeros, thereby mitigating the impact of sparsity^30^. In addition, MaAsLin2 uses a generalized linear model framework, allowing the inclusion of all our covariates. All results are FDR-corrected. We found a significant association only for item recognition performance (**Figure 4.A**). *Prevotella copri* showed a negative association with participants’ item recognition (**Table S3**): (coef= -1.32, qval=0.17). No taxa were associated with pattern completion or pattern separation performance. Given the sparse distribution of *P*.*copri*, we compared performance between carriers and non-carriers. We found that carriers of *P. copri* showed significantly worse item recognition performance (*P. copri* W=1253, p= 0.008 than non-carriers (**Figure 4.B**).

**Figure 4.**
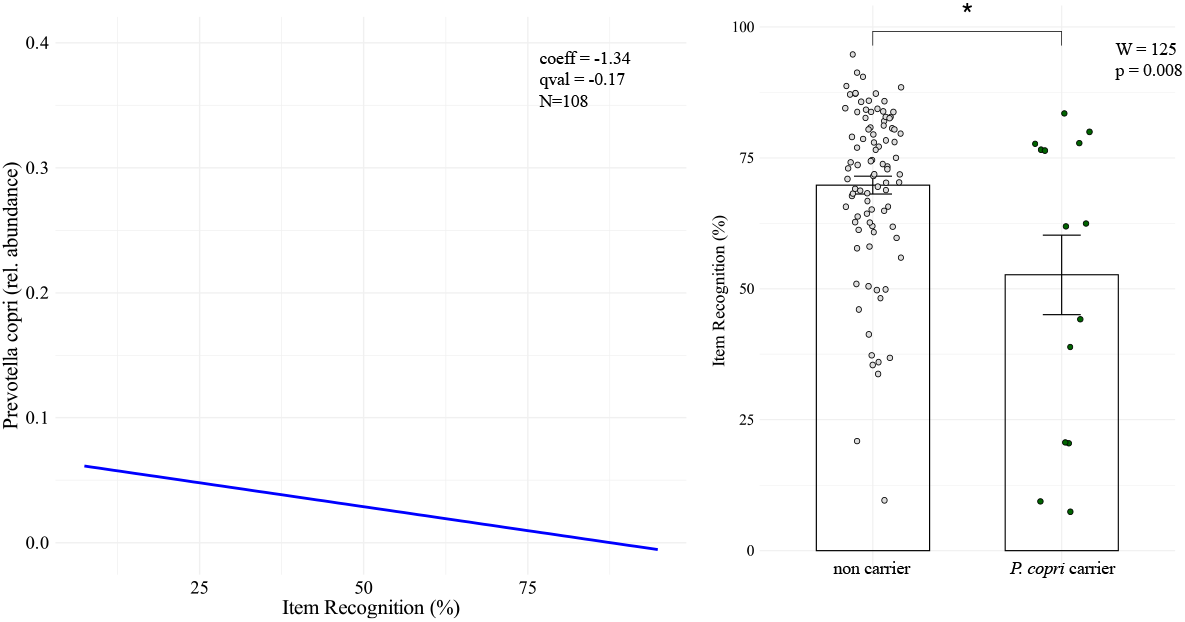
Association between *Prevotella copri* and Item Recognition Performance. (A) Scatter plot showing the negative association between *Prevotella copri* relative abundance and item recognition performance, as identified using a multivariable MaAsLin2 analysis (*coefficient* = -1.32, *qval* = 0.17). (B) Boxplot comparing item recognition performance between *Prevotella copri* carriers (participants with nonzero relative abundance) and non-carriers (participants with zero relative abundance). Wilcoxon rank-sum test results indicate that carriers exhibited significantly worse item recognition performance than non-carriers (*W* = 125, *p* = 0.008).

## DISCUSSION

Our results show that human episodic memory is associated with individuals’ indigenous - non-manipulated - gut microbiota composition. These results are consistent with previous findings using probiotics and prebiotics in rodents and a subset of similar studies in humans ^14,31^. Our results demonstrate that we can detect microbiota signals in healthy individuals related to how well participants performed an item memory recognition task. First, we found that greater alpha diversity was associated with better item recognition. Second, we observed a significant association between individuals’ item recognition performance and gut microbiota dissimilarity index (beta diversity). Thus indicating that the variations in performance among individuals were related to their gut microbiota structure. Finally, we found *Prevotella copri* significantly associated with item recognition. No associations were observed between pattern completion, pattern separation, and individuals’ gut microbiota composition.

The idea that low gut microbiota diversity is associated with disease has been reported in some studies. For example, compared to healthy individuals, low alpha diversity has been reported in patients with generalized anxiety^32^, depression^33^, Alzheimer’s disease^34^, Huntington’s disease^35^, and age-driven cognitive and physical decline^36,37^. In line with these studies, healthy adults with lower microbiota alpha diversity performed worse in item recognition. However, a previous study with an older Australian population (60-77 years old) did not find a relationship between individuals’ alpha diversity and their performance on any cognitive tasks, including episodic memory. One possibility is that they were underpowered to see this relationship, given their smaller sample size (n=69). Alternatively, differences in population age (younger in ours), the memory task employed (Mnemonic similarity task in our case and Quality of Episodic Secondary Memory from the Cognitive Drug Research battery in theirs), and different data processing methods might have contributed to different results.

More importantly, we found that the overall structure of the gut microbiota (as measured by beta diversity) was significantly associated with variability in item recognition. This is, variation in item recognition performance corresponded to variation in gut microbiota composition. Notably, the association between item recognition and gut microbiota structure between participants remained significant after controlling for several factors (sex, age, BMI, diet, gastrointestinal health, mode of birth, breastfeeding, and current pet cohabitation) widely reported to affect microbiota alpha and beta diversity. Additionally, the model showed that pet ownership during childhood and being breastfed were also associated with beta diversity, reinforcing their role in shaping the gut microbiota. However, these early-life factors were not directly associated with item recognition performance, suggesting that while they contribute to how similar microbiota composition is across individuals, their relationship to memory is not straightforward.

When looking for specific taxa related to memory, we found one species negatively associated with item recognition performance: *Prevotella copri*. Given the sparse distribution of *P*.*copri*, we compared item recognition performance between carriers and non-carriers and found that carriers of *P. copri* performed significantly worse than non-carriers (**Figure 4B**). Note that MaAsLin2 is a tool specifically designed for microbiome data analysis, providing robust results even for sparse and non-normally distributed taxa such as *P*.*copri*. However, given our dataset’s small sample size, these results warrant further validation in larger populations.

Although the concept of enterotypes has been debated^38^, *Prevotella copri* is one of the few species consistently forming a distinctive enterotype^39^ (a distinctive microbial community type in the human gut microbiota) in the human population. *Prevotella* has been associated with the metabolism of complex carbohydrates and plant-rich dietary patterns^40^. However, exploratory analyses showed that the Mediterranean diet was not associated with the relative abundance of *P*.*copri* (see **Table S4**). Nevertheless, we cannot rule out the possibility that *Prevotella copri* was associated with dietary habits that were not Mediterranean-related.

*Prevotella* has been associated with cognition in previous studies. A study comparing high and low *Prevotella* enterotypes in healthy women found that those with high *Prevotella* showed smaller hippocampal volume than low or non-*Prevotella* carriers^41^. Note, however, that in our dataset, we found effects at the species level for *Prevotella copri* but not at the genus level. In line with our findings, *Prevotella copri* in Alzheimer’s patients has been positively correlated with the disease’s progression, and it has been consistently linked to several autoimmune disorders of the central nervous system^40,42,43^. Nevertheless, different study results could be explained by genetic diversity among species^44^, differences in participants’ age, the study’s geographical location, or other confounding factors that have not been accounted for or controlled.

The finding that gut microbiota composition was only related to item recognition, but not pattern completion or pattern separation, suggests that the influence of gut microbiota composition on episodic memory may be widespread across hippocampal-cortical networks rather than targeting processes selective to very fine-grained mnemonic discrimination hypothesized to be reliant on specific hippocampal subfields. On the other hand, it might be the case that we were underpowered to see these effects. Studies in rodents using pathogenic bacteria or lipopolysaccharide^11,45^ injections that mimic the inflammatory effects of some gut bacteria have shown impairments in pattern separation performance. It is possible, then, that the influence of gut microbiota on pattern separation or pattern completion could only be seen under critical conditions (e.g., inflammation, aging, disease, developing) and not in healthy individuals.

In line with studies showing that gut microbiota modulates novel object recognition performance in rodents^2,4,6,7^, we found that the gut microbiota is related to how humans recognize items in a mnemonic similarity task. In addition, *Prevotella copri* and its associated gut microbiota composition appeared to play a key role in determining this relationship. Our findings set the path to investigating potential underlying mechanisms connecting gut microbiota composition and its influence on episodic memory. Although our data cannot speak of underlying mechanisms or directionality of effects, we can speculate on one potential path of action. Given previous links between *P. copri* and immune-related pathologies, these taxa may be associated with a microbiota composition that favors inflammation, negatively impacting hippocampal function, as shown by other studies^4^. Understanding how the gut microbiota can modulate episodic memory in its homeostatic – non-altered state – can open new targets for studying and potentially treating neurological (e.g., dementia, Alzheimer’s) and psychiatric pathologies (e.g., depression and anxiety). However, using large datasets and multi-omics will be fundamental to uncovering the mechanism behind the connection between gut microbiota and human episodic memory.

## METHODS

### Participants

One hundred and twenty-seven participants, 73 females and 54 men, aged 20 to 50 years (M = 26.55, SD = 7.66), and with a body mass index of 16.42 to 37.97 (M = 24.48, SD = 4.37) were recruited from the New York University (NYU) community via posted flyers and online cloud-based subject pool software (Sona Systems). The NYU Committee on Activities Involving Human Subjects approved all recruiting instruments and experimental protocols. Participants were remunerated $15 per hour and earned $50 for providing a stool sample. Eligible participants were proficient in English, had normal or corrected vision, were not currently taking psychoactive medications, corticosteroids, antibiotics, or probiotics, and were not diagnosed with psychiatric, neurological, or gastrointestinal disorders. Twelve participants were excluded for answering less than 80% of trials, two for not having a complete dataset, four for having a low-quality sample (with less than 10K read counts), and one for having a below-chance performance. Our final sample consisted of 108 participants (men=45, women=63), 18-50 years old (mean=25.59, std=6.89).

### Memory Task

We employed the standardized Mnemonic Similarity Task^46^. Participants underwent an incidental encoding session, where they were presented with 128 color images of everyday objects. Images were presented for 2 seconds with an intertrial interval of 0.5 seconds. Upon each image presentation, participants were instructed to indicate whether it depicted an ‘indoor’ or ‘outdoor’ object. After the incidental encoding, participants took a 25-minute break while completing questionnaires regarding covariates information (see Covariates below). After the break, participants underwent a surprise recognition memory test. Participants were presented with 64 *repetitive* items (same images shown in the previous session), 64 *foil* items (new objects not shown during the last session), and 64 *lure* items similar to those presented previously but not the same ones. Participants were instructed to indicate, upon each object presentation, whether it was ‘old’ (*repetitive*), ‘new’ (*foil*), or ‘similar’ (*lure*). We calculated *item recognition* as the number of ‘old’ responses to *repetitive* items (hits) minus the number of ‘old’ responses to *foil* items (errors). *Pattern completion* was calculated as the number of ‘old’ responses to *lure* items minus the number of ‘old’ responses to *foil* items. *Pattern separation* was calculated as the number of ‘similar’ responses to *lure* items (hits) minus the number of ‘similar’ responses to *foil* items (errors).

To ensure task engagement, we included only participants who responded to at least 80% of trials in the incidental encoding session in the analyses.

### Covariates

All analyses were controlled for factors that can shape gut microbiota communities^47^. Participants completed a series of demographic and psychological profiles questionnaires that we used to calculate the following covariates: age, sex, BMI, pet cohabitation during childhood (yes or no answer to the question *“Was there a pet in your family before you were three years old”)*, current pet cohabitation (yes or no question to *“do you currently live with a pet?”*), mode of birth *(“Were you born through natural birth or c-section?”*), breastfeeding (“*When you were a baby, where you breastfed or formula fed?”*), exercise (“*Do you frequently exercise during the week?”*), diet (participants answered the Mediterranean Diet Adherence Screener^48^), and gastrointestinal health (participants answered a gastrointestinal health questionnaire; see questionnaire in Supplementary Materials). We also included subclinical measures of depression, anxiety, and stress by employing the DASS 21^49^ questionnaire with its three subscales: Depression, Anxiety, and Stress scores.

### Stress reactivity; stress-induced cortisol responses

As part of the psychological profiling, participants performed a stress-induced reactivity test using the Cold Pressor Test^50,51^. Participants were asked to submerge their non-dominant hand and arm into ice-cold water (0°C - 3°C). Saliva samples were collected to measure basal cortisol levels and cortisol response to the stressor. We collected one saliva sample before the CPT (baseline) and two samples at 10 and 25 minutes after the CPT (see protocol in Figure S2). As an index of stress reactivity, we calculated the Area Under the Curve (AUC) using the trapezoidal rule: AUC = 0.5 * 15 * (baseline + 2 * t10 + t25, see distribution of stress reactivity values in FigureS3).

**Table 1.**
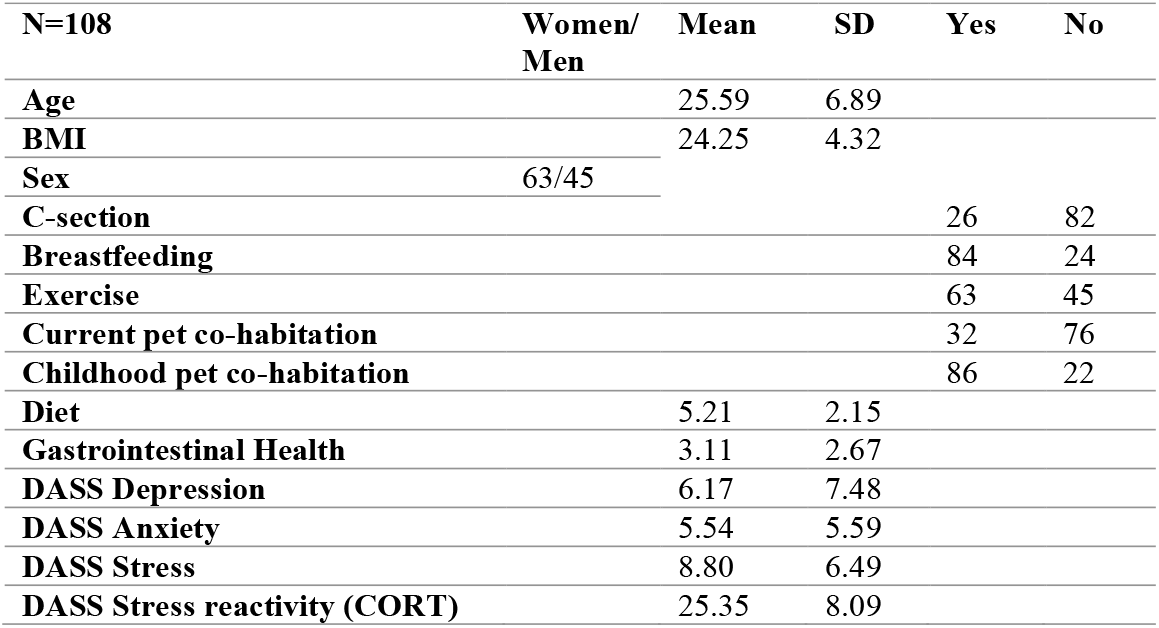
We included these variables as covariates in all models run to control for potential factors that could affect gut microbiome composition. Participants were asked; How old are you? How much do you weigh? How tall are you? What is your sex? Were you born through c-section? Were you breastfed? Do you frequently exercise during the week? Do you currently live with a pet? Did you have pets before you were 3 years old? Lastly, participants completed the Mediterranean Diet Adherence Screener, the DASS21 questionnaire, and a gastrointestinal health questionnaire (see gastrointestinal health questionnaire in Supplemental Results).

### Stool collection and DNA processing

At the end of the session, participants were provided with an OMR-200 OMNIgene-Gut stool collecting kit (DNA Genotek) and an OM-AC1 toilet accessory. Participants watched a video with instructions about using the collecting kit at home and received further verbal instructions from a research assistant who answered any questions. Participants were instructed to collect a stool sample during the next bowel movement and bring it to the lab the following day. Stool samples were stored at -20°C after receipt. Samples were processed and sequenced in one batch at the Columbia University Microbiome Core (CUMC). DNA was extracted using the MagAttract PowerSoil kit (Mo-Bio), and 16S rRNA amplicon sequencing of the V3-V4 region with the 341F/805R primer pair was performed on the Illumina MiSeq 2x300 platform following the manufacturer’s recommended best practices. The resulting FASTQ files were bioinformatically processed through the bioBakery AnADAMA2 0.90 workflows^52^. Sequences were demultiplexed with ea-utils 1.1.2 . Cutadapt 3.0^54^ was used to remove the remaining primer and adapter sequences. DADA2 1.14.1^55^ was used to denoise, filter, trim, merge, remove chimeras, and taxonomically classify the data, with default parameters other than a truncation length of 265bp for forward reads and 225bp for reverse as reads after these points have substantially lower average quality scores and interfered with read pair merging. Phylogenetic trees were constructed after the alignment of sequences using Clustal Omega 1.2.4 with FastTree2.1.10^57^. ASVs were taxonomically assigned using the SILVA 132^58^ reference database. Features were filtered, requiring at least 0.01% relative abundance in 10% of samples for all downstream analyses.

### Alpha diversity

Alpha diversity was quantified using the Inverse Simpson Index, which reflects the microbiota community’s richness (number of species/genus) and evenness (similarity of abundances of each species/genus). We performed linear regressions for each of the three memory measurements (item recognition, pattern separation, and pattern completion), including covariate factors (i.e., age, sex, BMI, mode of birth, breastfeeding, diet, exercise, and pet cohabitation during childhood and current).

### Beta diversity

Beta diversity was quantified by weighted Bray-Curtis distance. The variance explained (R^2^) and statistical significance for each memory measurement in multivariable models with control variables were calculated using Permutational Multivariable Analysis of Variance (PERMANOVA) tests. To visualize the community compositions, Bray-Curtis distances were ordinated by Principal Coordinates Analysis (PCoA, **Figure 2**). Calculations were performed with the vegan 2.5-7^59^ package, and visualizations were created using ggplot2 3.3.5 in the R 4.1.0 computing environment^60^.

### Feature associations (MaAsLin2)

To identify specific taxa associated with individuals’ memory measurements, we used general linear models implemented in MaAsLin2 1.73^61^. MaAsLin2 1.73 calculates each feature’s associations with metadata with multiple comparison adjustments across all models. We perform the analysis by grouping (glom) all features at the species level. If species-level annotations are missing, we group them at the highest known taxonomic rank or ‘terminal taxa. This technique allows dimensionality reduction and avoids several common analytic issues, such as very sparse data matrices (i.e., containing many zeros), overly stringent false discovery control, overfitting in random forest models, and a reduction in data noise.

Filtered, normalized relative abundance data is log-transformed before testing with a pseudo count of half of the lowest non-zero relative. Results were adjusted using the Benjamini and Hochberg method, and since this is an exploratory study, we reported as significant false discovery rate (FDR) with P-values of 0.25 or lower. We chose this cut-off as we believe it warrants not missing weaker but real associations, especially given the high dimensional nature of ‘omics data. Univariate and multivariate analyses were performed with 108 participants.

## DATA AVAILABILITY

*An OSF link with the behavioral and ASV data will be provided upon publication*.

## CODE AVAILABILITY

*A GitHub link will be provided upon publication with the scripts used to analyze the data*.

## ACKNOWLEDGEMENTS

We thank Dr. Anne-Catrin Uhlemann and Felix Rozenberg for their work on stool sample processing and sequencing. We are also grateful to Yasmine Elasmar and Eugenia Zhukovsky for their assistance with data collection. This research was supported by The Vulnerable Brain Project (vbp.life) awarded to JEL and EAP; the James S. McDonnell Foundation, NIH grant R21MH134157 and the Hodgson Fund from the Department of Psychology at Harvard University to EAP; and the NIH grant TL1TR001447 from the National Center for Advancing Translational Sciences (NCATS) to JPO.

## AUTHOR CONTRIBUTIONS

Conception and design; JPO, LD, EAP, JEL, Data acquisition; JPO, Data analysis; JPO, TK, Data interpretation; JPO, TK, XM, LD, EAP, CH, Writing Paper: JPO, TK, XM, LD, EAP, CH.

