## Supplementary Material for "Item recognition is associated with gut microbiota composition in healthy humans"

### 1. Taxonomic Profiling

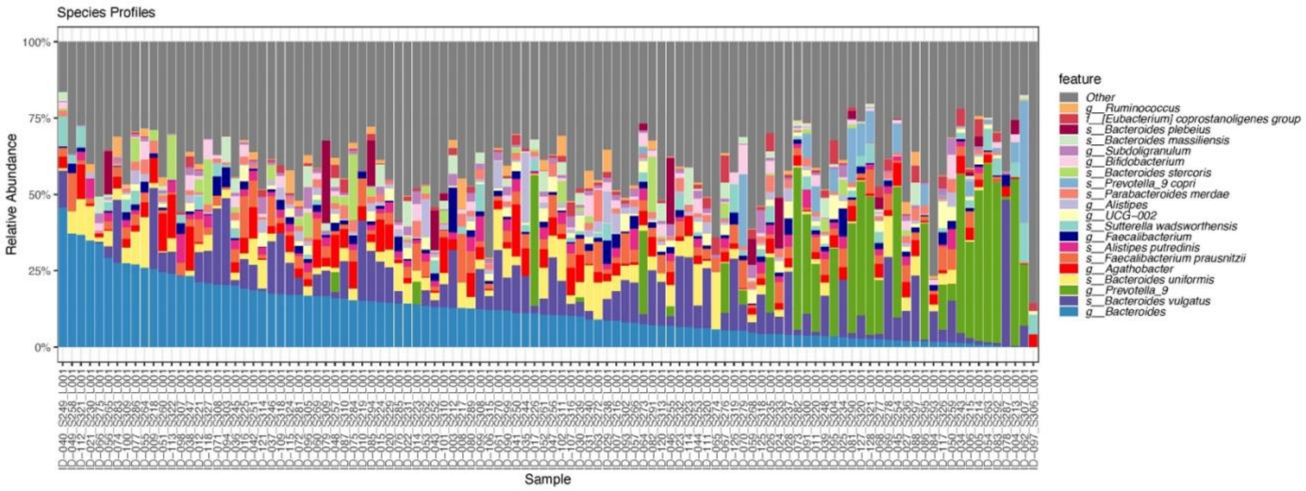


**Figure S1.** Relative Abundances of the top 20 most abundant species in the sample

### 2. Results for Multivariable ANOVA on Inverse Simpson Index

| | Sum Sq | Df | F value | Pr(>F) |
|---|---|---|---|---|
| **Item recognition** | **129.775** | **1** | **4.048** | **0.047** |
| Age | 4.001 | 1 | 0.124 | 0.724 |
| Sex | 50.938 | 1 | 1.588 | 0.210 |
| BMI | 11.952 | 1 | 0.372 | 0.543 |
| Mediterranean diet | 7.034 | 1 | 0.219 | 0.640 |
| Gastro health | 43.758 | 1 | 1.364 | 0.245 |
| Exercise frequency | 74.026 | 1 | 2.309 | 0.132 |
| **Breastfeeding** | **273.797** | **1** | **8.540** | **0.004** |
| C-section | 14.301 | 1 | 0.446 | 0.505 |
| Current pet | 26.003 | 1 | 0.811 | 0.370 |
| Childhood pet | 3.524 | 1 | 0.109 | 0.741 |
| DASS Depression | 3.618 | 1 | 0.112 | 0.737 |
| DASS Anxiety | 0.003 | 1 | 0.000 | 0.991 |
| DASS Stress | 0.390 | 1 | 0.012 | 0.912 |
| Stress reactivity (CORT) | 3.230 | 1 | 0.100 | 0.751 |
| Residuals | 2757.075 | 86 | NA | NA |

**Table S1A**. Multivariable ANOVA for Item Recognition on Inverse Simpson Diversity.

| | Sum Sq | Df | F value | Pr(>F) |
|---|---|---|---|---|
| **Pattern completion** | **1.739** | **1** | **0.051** | **0.820** |
| Age | 0.261 | 1 | 0.007 | 0.929 |
| sex | 38.809 | 1 | 1.156 | 0.285 |
| BMI | 11.142 | 1 | 0.332 | 0.565 |
| Mediterranean diet | 12.455 | 1 | 0.371 | 0.543 |
| Gastro Health | 23.633 | 1 | 0.704 | 0.403 |
| Exercise frequency | 85.699 | 1 | 2.554 | 0.113 |
| **Breastfeeding** | 282.607 | 1 | 8.424 | 0.004 |
| C-section | 12.217 | 1 | 0.364 | 0.547 |
| Current pet | 11.813 | 1 | 0.352 | 0.554 |
| Childhood pet | 1.416 | 1 | 0.042 | 0.837 |
| DASS Depression | 3.213 | 1 | 0.095 | 0.757 |
| DASS Anxiety | 4.056 | 1 | 0.120 | 0.728 |
| DASS Stress | 0.540 | 1 | 0.016 | 0.899 |
| Stress reactivity (CORT) | 7.558 | 1 | 0.225 | 0.636 |
| Residuals | 2885.111 | 86 | NA | NA |

**Table S1B.** Multivariable ANOVA for Pattern Completion on Inverse Simpson Diversity.

| | Sum Sq | Df | F value | Pr(>F) |
|---|---|---|---|---|
| **Pattern separation** | **52.277** | **1** | **1.586** | **0.211** |
| Age | 2.155 | 1 | 0.065 | 0.798 |
| BMI | 7.683 | 1 | 0.233 | 0.630 |
| Mediterranean diet | 8.070 | 1 | 0.244 | 0.621 |
| Gastro Health | 26.780 | 1 | 0.812 | 0.369 |
| Exercise frequency | 67.151 | 1 | 2.037 | 0.157 |
| **Breastfeeding** | 260.741 | 1 | 7.910 | 0.006 |
| C-section | 6.946 | 1 | 0.210 | 0.647 |
| Current pet | 10.836 | 1 | 0.328 | 0.567 |
| Childhood pet | 1.614 | 1 | 0.048 | 0.825 |
| sex | 41.054 | 1 | 1.245 | 0.267 |
| DASS Depression | 7.157 | 1 | 0.217 | 0.642 |
| DASS Anxiety | 10.613 | 1 | 0.322 | 0.571 |
| DASS Stress | 0.613 | 1 | 0.018 | 0.891 |
| Stress reactivity (CORT) | 7.206 | 1 | 0.218 | 0.641 |
| Residuals | 2834.573 | 86 | NA | NA |

**Table S1C.** Multivariable ANOVA for Pattern Separation on Inverse Simpson Diversity.

### 3. Multivariable PERMANOVA on Bray-Curtis Dissimilarity

| | Df | SumOfSqs | R2 | F | Pr(>F) |
|---|---|---|---|---|---|
| **Item recognition** | **1** | **0.395** | **0.017** | **1.866** | **0.037** |
| age | 1 | 0.162 | 0.007 | 0.765 | 0.726 |
| sex | 1 | 0.214 | 0.009 | 1.012 | 0.394 |
| BMI | 1 | 0.184 | 0.008 | 0.869 | 0.607 |
| Mediterranean diet | 1 | 0.315 | 0.014 | 1.489 | 0.115 |
| Gastro health | 1 | 0.195 | 0.008 | 0.921 | 0.518 |
| Exercise frequency | 1 | 0.241 | 0.010 | 1.140 | 0.285 |
| **Breastfeeding** | **1** | **0.376** | **0.017** | **1.778** | **0.044** |
| C-section | 1 | 0.181 | 0.008 | 0.855 | 0.613 |
| Current pet | 1 | 0.152 | 0.006 | 0.719 | 0.782 |
| **Childhood pet** | **1** | **0.456** | **0.020** | **2.154** | **0.015** |
| DASS Depression | 1 | 0.327 | 0.014 | 1.547 | 0.086 |
| DASS Anxiety | 1 | 0.167 | 0.007 | 0.790 | 0.649 |
| DASS Stress | 1 | 0.197 | 0.008 | 0.931 | 0.475 |
| Stress reactivity (CORT) | 1 | 0.165 | 0.007 | 0.779 | 0.712 |
| Residual | 86 | 18.222 | 0.822 | NA | NA |
| Total | 101 | 22.146 | 1 | NA | NA |

**Table S2A.** Multivariable PERMANOVA for Item recognition on Bray-Curtis dissimilarity at the species level, relative abundance with 999 permutations.

| | Df | SumOfSqs | R2 | F | Pr(>F) |
|---|---|---|---|---|---|
| Pattern Completion | 1 | 0.204 | 0.009 | 0.973 | 0.436 |
| Age | 1 | 0.216 | 0.009 | 1.032 | 0.386 |
| BMI | 1 | 0.186 | 0.008 | 0.889 | 0.580 |
| Mediterranean diet | 1 | 0.326 | 0.015 | 1.558 | 0.108 |
| Gastro Health | 1 | 0.227 | 0.010 | 1.085 | 0.322 |
| Exercise frequency | 1 | 0.217 | 0.010 | 1.035 | 0.381 |
| Creastfeeding | 1 | 0.359 | 0.016 | 1.716 | 0.066 |
| C-section | 1 | 0.173 | 0.008 | 0.829 | 0.643 |
| Current pet | 1 | 0.128 | 0.005 | 0.612 | 0.898 |
| **Childhood pet** | **1** | **0.439** | **0.020** | **2.097** | **0.019** |
| Sex | 1 | 0.203 | 0.009 | 0.968 | 0.436 |
| DASS Depression | 1 | 0.327 | 0.015 | 1.559 | 0.090 |
| DASS Anxiety | 1 | 0.204 | 0.009 | 0.976 | 0.431 |
| DASS Stress | 1 | 0.191 | 0.008 | 0.911 | 0.503 |
| Stress Reactivity (CORT) | 1 | 0.171 | 0.007 | 0.819 | 0.640 |
| Residual | 86 | 18.033 | 0.831 | NA | NA |
| Total | 101 | 21.698 | 1 | NA | NA |

**Table S2B.** Multivariable PERMANOVA for Pattern Completion on Bray-Curtis Dissimilarity at genus level relative abundance with 999 permutations.

| | Df | SumOfSqs | R2 | F | Pr(>F) |
|---|---|---|---|---|---|
| Pattern separation | 1 | 0.205 | 0.009 | 0.982 | 0.411 |
| Age | 1 | 0.222 | 0.010 | 1.060 | 0.346 |
| sex | 1 | 0.202 | 0.009 | 0.966 | 0.434 |
| BMI | 1 | 0.191 | 0.008 | 0.911 | 0.553 |
| Mediterranean diet | 1 | 0.310 | 0.014 | 1.481 | 0.123 |
| Gastro Health | 1 | 0.223 | 0.010 | 1.067 | 0.342 |
| Exercise frequency | 1 | 0.214 | 0.009 | 1.023 | 0.401 |
| breastfeeding | 1 | 0.357 | 0.016 | 1.704 | 0.066 |
| C-section | 1 | 0.144 | 0.006 | 0.690 | 0.818 |
| Current pet | 1 | 0.121 | 0.005 | 0.581 | 0.920 |
| **Childhood pet** | **1** | **0.435** | **0.020** | **2.074** | **0.021** |
| DASS Depression | 1 | 0.318 | 0.014 | 1.5174 | 0.097 |
| DASS Anxiety | 1 | 0.216 | 0.009 | 1.034 | 0.366 |
| DASS Stress | 1 | 0.190 | 0.008 | 0.908 | 0.506 |
| Stress reactivity (CORT) | 1 | 0.172 | 0.007 | 0.820 | 0.641 |
| Residual | 86 | 18.031 | 0.831 | NA | NA |
| Total | 101 | 21.698 | 1 | NA | NA |

**Table S2C.** Multivariable PERMANOVA for pattern Separation on Bray-Curtis at genus level relative abundance with 999 permutations.

### 4. Significant Single Taxa Associations for the Item Recognition Multivariable Model

| feature | feature | coef | stderr | N | N.not.0 | pval | qval |
|---|---|---|---|---|---|---|---|
| s_*Prevotella 9 copri* | Item recognition | -1.321 | 0.336 | 108 | 21 | <0.001 | 0.177 |
| f_Coriobacteriales incertae sedis | Sex | -1.563 | 0.389 | 108 | 29 | <0.001 | 0.177 |
| o_DTU014 | Breastfeeding | -1.134 | 0.280 | 108 | 24 | <0.001 | 0.177 |

**Table S3.** Single feature significant associations were found with the multivariable model for item recognition (MaAsLin2, multivariable model).

### 5. Exploratory analysis for Mediterranean diet.

| feature | coef | stderr | N | N.not.0 | pval | qval |
|---|---|---|---|---|---|---|
| s__Dielma.fastidiosa | 0.171 | 0.049 | 108 | 20 | <0.001 | 0.164 |
| g__Lachnospira | 0.457 | 0.142 | 108 | 46 | 0.001 | 0.183 |

**Table S4.** Single feature significant associations with the Mediterranean diet **(**MaAsLin2, univariable model)

Although *P. copri* was not associated with the Mediterranean diet, caution is advised when using self-reported dietary questionnaires, as they can be subject to recall bias and memory inaccuracies. Nevertheless, the Mediterranean Diet Adherence Score (MEDAS) used in this study is a widely accepted and validated tool in nutritional epidemiology, providing a practical and reliable measure of dietary habits[1–3]. Nevertheless, future studies could strengthen dietary assessments by incorporating objective biomarkers or detailed food frequency questionnaires (FFQs) to complement self-reported data.

### 6. Stress reactivity protocol

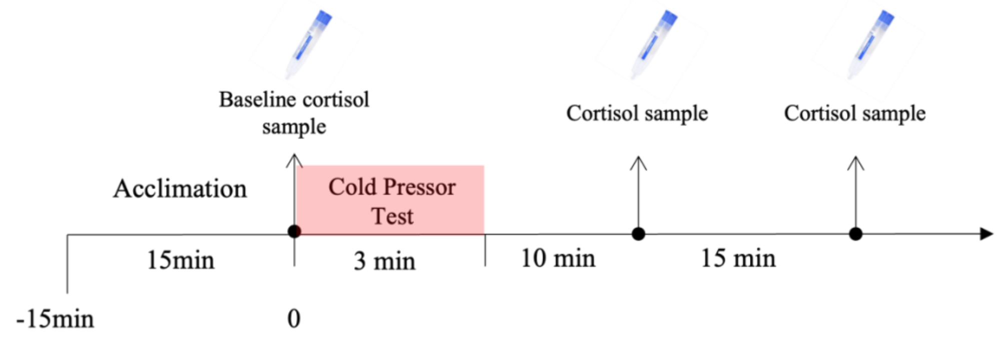


**Figure S2.** Cold Pressor Test protocol. The day after the Memory tasks, participants underwent The Cold Pressor Test (CPT)[4,5], which was used to provoke physical stress. Participants were asked to submerge their non-dominant hand and arm into ice-cold water (0°C - 3°C). Saliva samples were collected to measure basal cortisol levels and cortisol response to the stressor. We collected one saliva sample before the CPT (baseline) and two samples at 10 and 25 minutes after the CPT. Right after the CPT, participants rated their level of unpleasantness on a scale from 1 to 10 (1 “not at all unpleasant” to 10 ”very much unpleasant”). On average, participants rated their experience as very unpleasant (average 8).

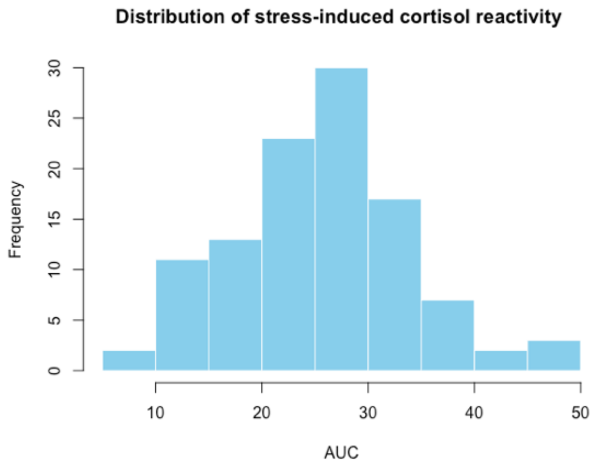


**Figure S3. Distribution of stress reactivity measure by cortisol stressed-induce cortisol release.** As an index of stress reactivity, we calculated the area under the curve between the three cortisol measures using the trapezoidal rule. AUC = 0.5 * 15 * (baseline cort + 2 * t10cort + t25cort). *AUC= area under the curve.

### 7. Gastrointestinal Health Questionnaire

1. Have you been diagnosed with a functional gastrointestinal disorder (e.g., irritable bowel syndrome, Crohn's disease)? If so, please specify diagnosis and age of diagnosis. -> *Participants were discarded if they answered 'Yes' to this question.*

2. Symptoms in the upper abdomen

A. In the last two months, how often did you have pain or an uncomfortable feeling in the upper abdomen (above the belly button)? PLEASE TICK THE CORRECT RESPONSE.
a. Never ___
b. 1 to 3 times per month ___
c. Once a week ___
d. Several times a week ___
e. Every day ___

B. Which of the following feelings did you have above the belly button? PLEASE TICK THE CORRECT RESPONSE.
a. Pain ___
b. Nausea ___
c. Bloating ___
d. Feeling of fullness ___
e. Not being hungry after eating very little ___

3. Symptoms Lower abdomen

C. In the last two months, how often did you have pain or an uncomfortable feeling in the lower abdomen (around or below the belly button)? PLEASE TICK THE CORRECT RESPONSE.
a. Never ___
b. 1 to 3 times per month ___
c. Once a week ___
d. Several times a week ___
e. Every day ___

D. Which of the following feelings did you have below the belly button?
a. Pain ___
b. Nausea ___
c. Bloating ___
d. Feeling of fullness ___
e. Not being hungry after eating very little ___

4. In the last two months, how often did you have diarrhea? PLEASE TICK NEXT TO THE CORRECT RESPONSE.
a. Never ___
a. 1 to 3 times per month ___
b. Once a week ___

c. Several times a week ___
d. Every day ___

5. In the last two months how often did you experience a feeling of bloating? PLEASE TICK NEXT TO THE CORRECT RESPONSE.
a. Never ___
b. 1 to 3 times per month ___
c. Once a week ___
d. Several times a week ___
e. Every day ___

6. In the last two months, how often did you have constipation with compensatory diarrhea? PLEASE TICK NEXT TO THE CORRECT RESPONSE.
a. Never ___
b. 1 to 3 times per month ___
c. Once a week ___
d. Several times a week ___
e. Every day ___

7. In the last two months, how often did you have constipation without compensatory diarrhea? PLEASE TICK NEXT TO THE CORRECT RESPONSE.
a. Never ___
b. 1 to 3 times per month ___
c. Once a week ___
d. Several times a week ___
e. Every day ___
